# A Yeast Logic Toolkit for Fast and Efficient Assembly of Genetic Circuits

**DOI:** 10.64898/2026.09.21.753091

**Authors:** Maik Molderings, Erik Kubaczka, Heinz Koeppl

## Abstract

Synthetic biologists aim for standardization and simplicity in order to ensure reproducibility and reliability. Toolkits offer both, making it easier to share DNA sequences between labs while minimizing context dependent effects from sequences surrounding the construct of interest. The Yeast Toolkit (YTK) for the model organism *Saccharomyces cerevisiae* has found wide adoption within the scientific community. While the Toolkit and its existing extensions offer a large variety of different parts, there is a lack of characterized logic gates and insulator containing backbones for the assembly of genetic circuits with the yeast toolkit syntax. In this work, we present the Yeast Logic Toolkit (YLT). YLT contains 40 pre-assembled backbones with dropouts, 109 parts for NOT/NOR gate assembly, and a subdivided promoter Type 2 part that allows more flexible 5’UTR design. For insulation, the backbones contain spacer sequences and self-cleaving ribozymes which buffer against transcriptional read-trough from upstream sequences. Due to our subdivision of Type 2 parts, we are able to manipulate the 5’ untranslated region without a cloning scar upstream of the start codon. We also characterized the NOT and NOR gates from the Cello library for yeast in the sequence context of the yeast toolkit cloning scheme. This toolkit extension will make it easier and faster for scientists to assemble genetic circuits in yeast while providing a minimum of context dependent effects such as genetic read-trough due to the standardization of up- and downstream sequences surrounding the transcriptional units.

**TOC Graphic:** 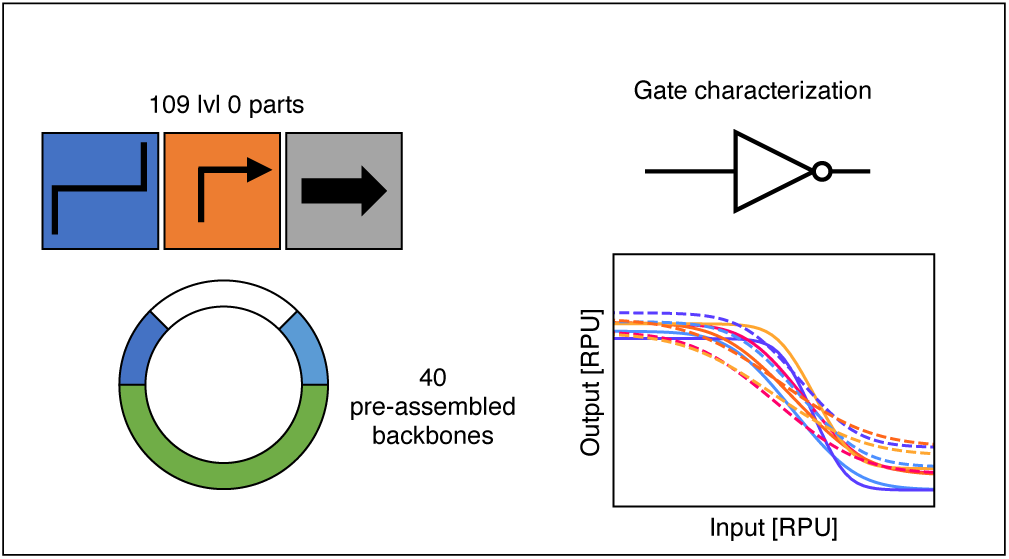

## 1 Introduction

Synthetic biology, a discipline which tries to apply engineering principles to life sciences, aims for reproducibility, which is an inherent necessity of engineering. One building block for reproducibility is standardization (*1*). In order to standardize cloning, which is the basis for many biologists in the lab to tackle biological questions, many techniques have been developed which allow for standardization. One of them is Golden Gate cloning, which is a one-pot reaction using Type IIS restriction enzymes allowing researchers to standardize cloning via usage of arbitrary overhangs (*2*). In the following years, Golden Gate-based cloning became more and more popular. In the budding yeast *S. cerevisiae* one of the most used techniques is the yeast toolkit (YTK), which is based on modular cloning (MoClo), a Golden Gate-based cloning scheme (*3*, *4*). Since then, many extensions have been published, offering easier strain engineering (*5*), genome editing via CRISPR (*6*, *7*), optogenetics (*8*), protein secretion and surface display (*9*), intracellular protein targeting (*10*), and inducible promoters (*11*).

Despite these advances, the design and implementation of reliable synthetic regulatory circuits in yeast remains a significant challenge. Yeast combines advantages of a well-characterized eukaryotic chassis with industrial relevance, making it attractive for studying gene regulation and building sophisticated control systems (*12*). However, circuit prototyping requires that individual gates respond consistently across different circuit contexts, and that transcriptional units are insulated from unwanted interactions. These requirements go beyond what current YTK-based kits provide, as their focus lies in strain engineering and application-specific modules rather than systematic circuit construction.

Initial progress has been made toward standardized circuit design. One of the most prominent efforts is Cellular Logic (Cello) (*13*), which was first implemented in *E. coli* and later extended to yeast (*14*). The yeast Cello library contains a set of bacterial repressors and synthetic promoters realizing NOT and NOR gates. To harmonize gate performance, synthetic 10 bp Kozak sequences (*14*, *15*) have been used to adjust translation initiation rates, producing similar response functions. Furthermore, insulation strategies such as self-cleaving ribozymes and terminators have been developed to reduce transcriptional read-through and crosstalk between transcriptional units (*16–18*). While these approaches demonstrate the feasibility of genetic logic in yeast, they have not yet been integrated into a YTK-compatible cloning framework that enables rapid and standardized circuit assembly.

In this work, we developed the Yeast Logic Toolkit (YLT), a YTK-compatible toolkit for the assembly of insulated synthetic circuits in yeast. YLT incorporates promoters, repressors, and insulator elements derived from the Cello library (*14*), while remaining compatible with the YTK toolkit (*4*) and the MYT extension for larger numbers of transcriptional units and easier strain engineering (*5*). To increase design flexibility, we further subdivided Type 2 parts to enable customizable 5’ untranslated region (5’ UTR) design. We then characterized the performance of the resulting gates and insulators in the context of the toolkit and demonstrated its functionality within the YTK framework.

## 2 Results

### 2.1 Adaptation of the Cello library into the YTK framework

To establish YLT, we adapted gate components of the Cello library to the YTK cloning framework and introduced changes that support insulation between adjacent transcriptional units, as well as more freedom for the 5’ UTR design. YLT is designed as a bridge between the Cello gate library and YTK-derived cloning systems (Figure 1 a). Insulator sequences were embedded directly in the type 1 connectors and used for the assembly of 40 pre-assembled level 1 backbones (Figure 1 b), while the original type 2 was separated into 2a and 2b parts in order to separate the promoter and the 5’ UTR, allowing its flexible design within the standard hierarchical assembly scheme (Figure 1 c,d).

**Figure 1:**
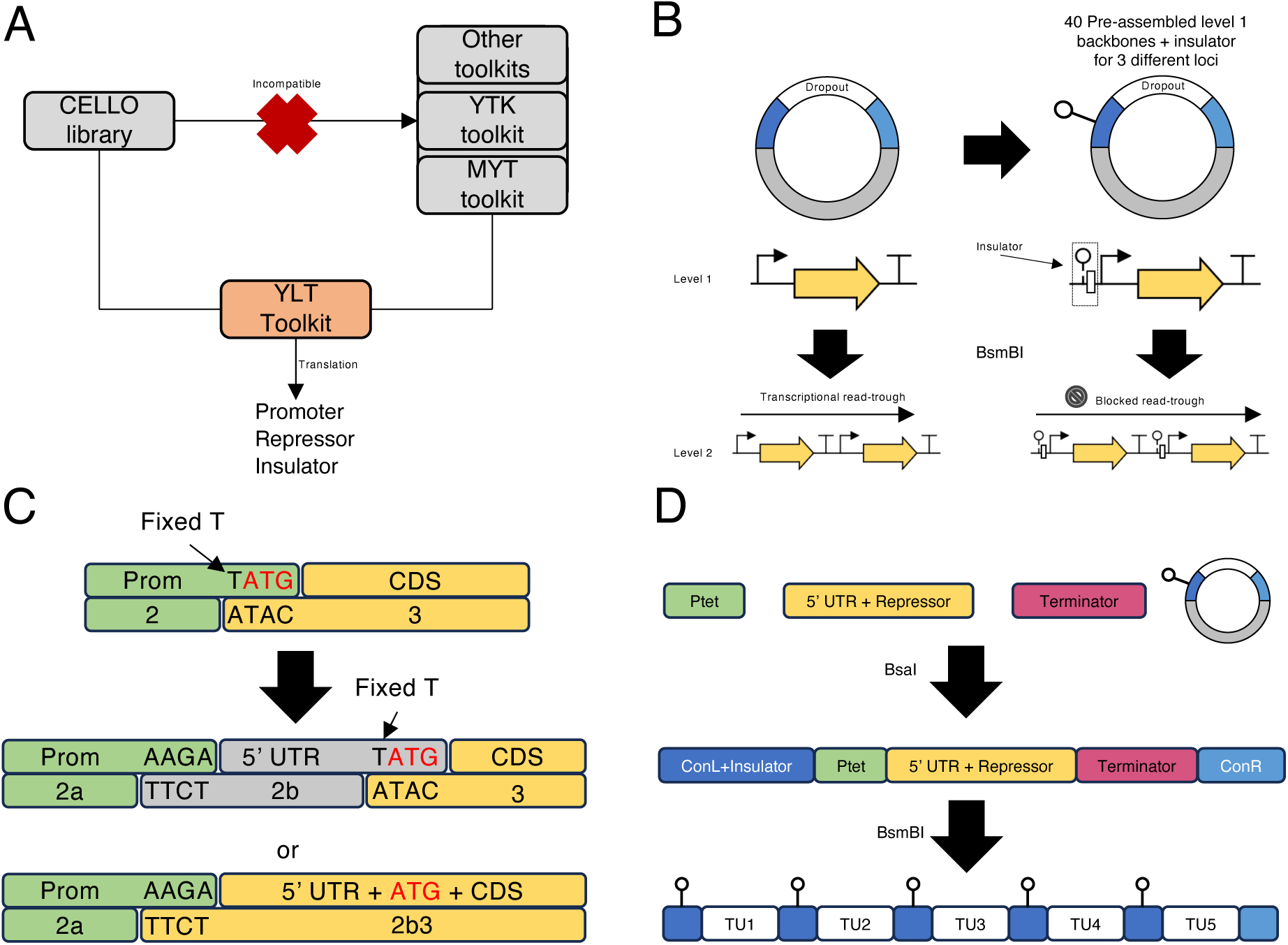
Architecture of the YLT toolkit. **(A)** YLT bridges the incompatibility between the Cello yeast library and MoClo-based yeast toolkits such as YTK and MYT by reformatting Cello-derived parts into a compatible assembly framework. **(B)** YLT includes 40 pre-assembled Level 1 backbones for three different loci, each containing insulator sequences within the Type 1 connector regions. These insulators consist of a ribozyme and spacer sequence and are intended to reduce transcriptional readthrough from an upstream transcriptional unit by ribozyme self-cleavage when transcription does not terminate efficiently. **(C)** In the YTK toolkit, the junction between promoter and CDS uses the TATG overhang, resulting in a fixed T at position -1 relative to the start codon. YLT introduces a separate Type 2b part for the 5’ UTR, or alternatively a combined Type 2b3 part, thereby enabling more flexible 5’ UTR design and removal of this constraint. **(D)** Within the hierarchical MoClo workflow, YLT enables assembly of Level 1 transcriptional units with customized 5’ UTR/CDS combinations, while insulated Level 2 constructs are generated through the use of the YLT Level 1 backbones.

A central design feature of YLT is the inclusion of insulator sequences. We redesigned type 1 connectors and included self-cleaving ribozymes (*16–18*), as well as spacer sequences. By using these redesigned type 1 connectors, we constructed 40 level 1 backbones (Figure 2) and Supporting Table 1) with dropout cassettes already containing the insulator sequences in the suggested order (*14*). When assembling a level 2 plasmid, the insulators are automatically placed between the transcriptional units (Figure 1 d). The backbones can be used for three different integration sites and are fully compatible with both YTK and MYT. Therefore, YLT can be integrated into existing yeast cloning workflows without requiring a separate assembly standard.

**Figure 2:**
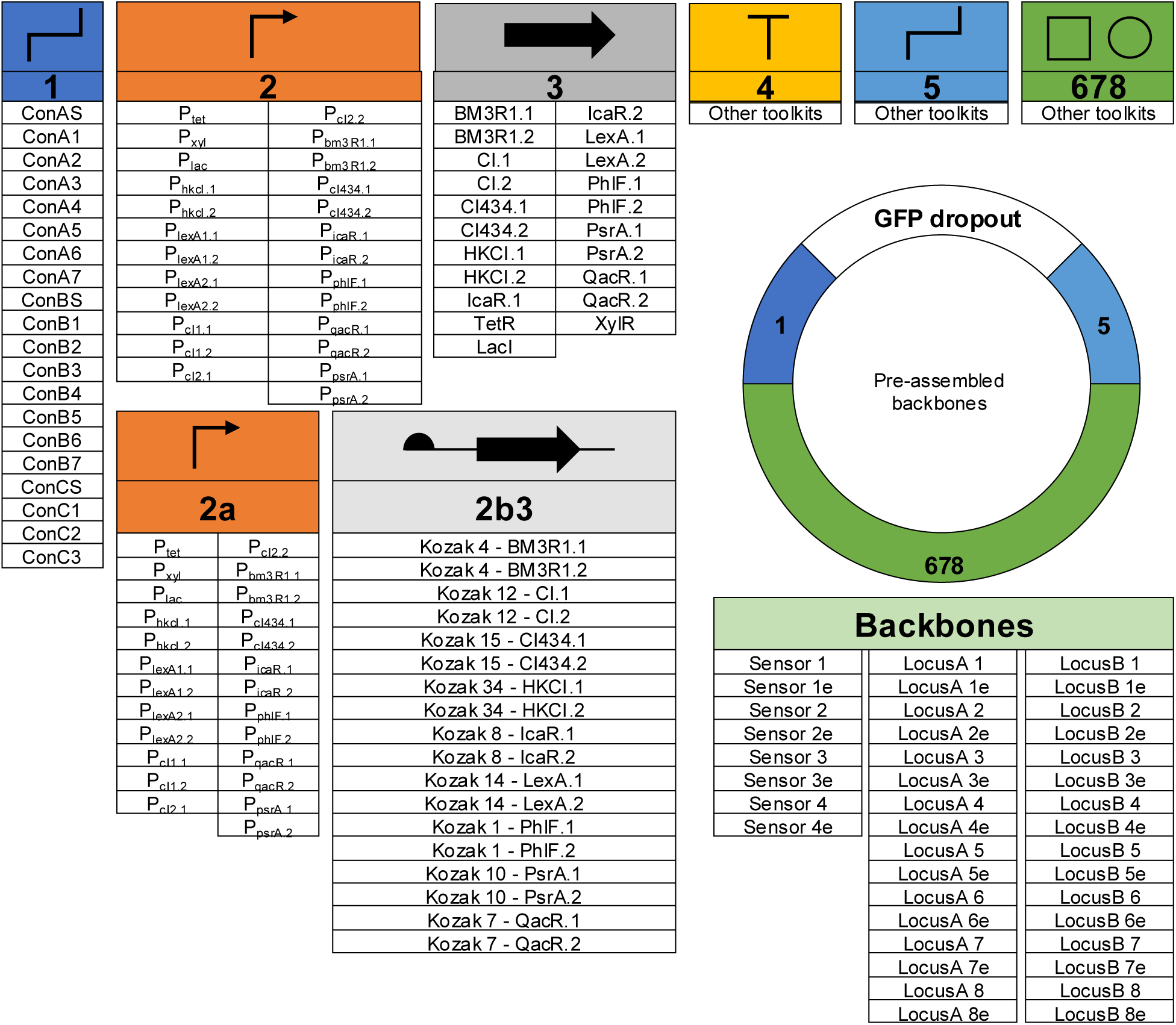
Overview of parts in the YLT toolkit. Level 1 plasmids can be assembled by combining parts that span number 1-8. Backbones act as a part combination with a GFP dropout which can be replaced by parts spanning number 2-4.

The toolkit also provides all promoter and repressor coding sequences as standard, unmodified type 2 and type 3 parts (Figure 2). This allows users to repurpose these parts outside the context of standardized gates and circuits, using the conventional YTK Type 2–Type 3 assembly grammar for applications unrelated to standardized circuit construction. Both formats use the same sequences and can be freely combined within the same assembly. Therefore, users can choose the assembly format that best fits their intended application.

In order to achieve comparable response functions, repressors in the Cello library were fused with synthetic Kozak sequences of different strength (*14*, *15*). Because the Kozak sequences used do not have a T at position -1, we split the type 2 into subparts in order to design the 5’ UTR without the need for a TATG as overhang between promoter and CDS, which fixes the T at position -1. Therefore, we combine 5’ UTR design flexibility with YTK compatibility.

### 2.2 Kozak variant expression and insulator function in the YLT framework

While AAAA as an overhang between type 2a and 2b would have allowed for scarless assembly between Cello promoters, which all have AAAA at their 3’ end, AAAA is a suboptimal overhang in terms of ligation fidelity. We therefore chose AAGA as an overhang, which was predicted to be a high-efficiency overhang by the NEBridge Get-Set Tool (*19*). This decision reflects the fact that the YLT is not only designed for the assembly of synthetic circuits with the Cello library, but should also remain open to projects in which the promoters used do not end in AAAA.

Next, we tested the influence of the new AAGA overhang on synthetic Kozak sequences with known relative expression levels (*15*). Due to the overhang, the 5’UTR always starts with AAGA, which might change expression levels compared to the original 10 bp sequence. To test this, we generated a Kozak variant library at the Level 0 stage: parts are usually made by PCR with BsmBI and BsaI restriction sites in the primer sequence, and for genes with varying Kozaks, the Kozak sequence itself is included in the primer, yielding 2b3 parts with different Kozaks upstream of the gene’s start codon (Figure 3 a). Expression level are converted into relative promoter units (RPU) by subtracting background fluorescence (wildtype) and dividing by the P_Pfy1_ expression (*14*). Kozaks are ordered from left to right according to previously reported strength rankings (*15*). Expression levels do not simply reproduce the original ranking. Several low- and mid-strength Kozaks (e.g. Kozak 84) show expression levels that no longer follow their expected position, indicating that the AAGA overhang measurably alters the relative strength of individual Kozak sequences. However, the four Kozaks originally predicted to be strongest (Kozak 818, 1165, 1847, and 2042) remain among the four highest-expressing constructs with the AAGA overhang as well, suggesting that while absolute rank order is not preserved, the strongest Kozaks largely retain their relative advantage even in the AAGA context (Figure 3a).

**Figure 3:**
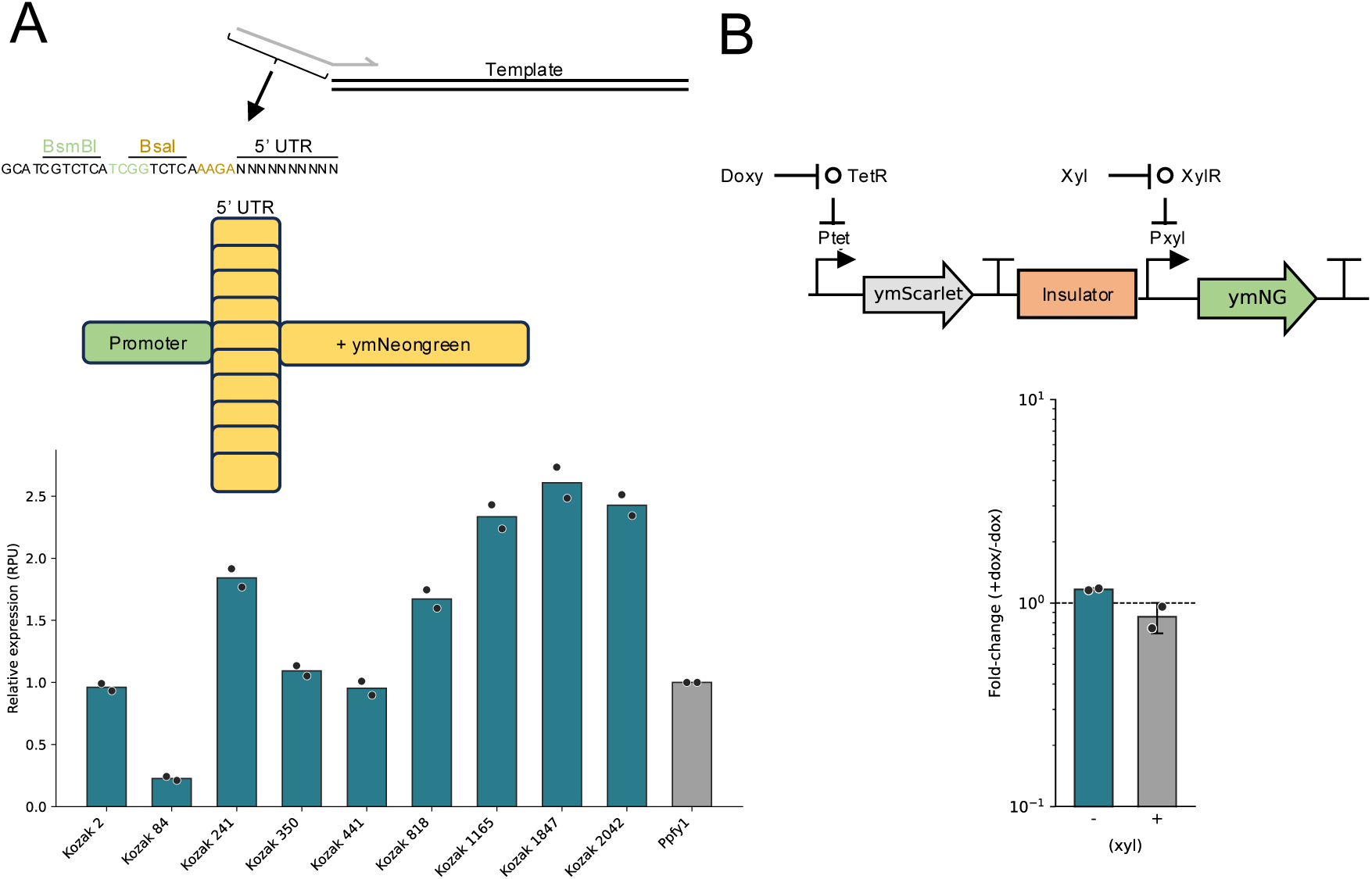
*A)* In order to design type 2b3 parts, a template CDS can be amplified via PCR by adding the 5’ UTR and the 2b overhang withing the sequence of the oligo. 10 Kozak sequences of different strength upstream of ymNeongreen were quantified via flow cytometry. *B)* Insulator verification of the different backbones from the YLT toolkit. If the insulator works, expression of ymNeongreen should be independent of the expression level of an upstream transcriptional unit. ymScarlet is only used as spacer and was not quantified.

Next, we tested whether the insulators incorporated into the type 1 parts remain functional when they are a fixed part of the level 1 backbones. In order to not interfere with BsmBI and BsaI reactions, we had to mutate existing restriction sites within the spacers, as well as change the RiboJ60 ribozyme (*14*, *16*, *20*) to a VtmoJ ribozyme (*16*, *21*). Similarly to (*14*), we tested insulator functionality by placing an insulator between two transcriptional units, an upstream unit with an inducible promoter driving ymScarlet and a downstream unit expressing ymNeongreen, and measuring ymNeongreen expression as a readout of insulation (Figure 3b). The fold-change in ymNeongreen expression with and without induction of the upstream expression unit remained close to one across both xylose conditions, indicating minimal transcriptional read-through from the upstream unit and confirming that the VtmoJ-based insulator retains its function.

### 2.3 Characterization of single-input sensors and NOT gates

To rule out locus-specific effects on the behavior of sensors and gates from the Cello library, we characterized them at the Int. 1, Int. 2, and Int. 3 loci of the MYT toolkit (*5*). To convert absolute fluorescence units into RPU, we measured the fluorescence of wildtype cells and of a strain containing the reference promoter P_Pfy1_ (Figure 4a) alongside each sensor strain. RPU values were then calculated by subtracting the wildtype background fluorescence and dividing by the background-subtracted fluorescence of the P_Pfy1_ reference strain. The xylose repressor (XylR), tetracycline repressor (TetR), and lactose repressor (LacI) were genetically integrated into the Int. 1 locus under constitutive promoters (*5*), while the corresponding inducible promoters P_xyl_, P_tet_, and P_lac_ were integrated into Int. 2, driving ymNeongreen expression. Flow cytometry measurements showed increasing ymNeongreen expression (Figure 4b) with increasing inducer concentration (xylose, doxycycline, and IPTG, respectively), as expected. Having confirmed functional single-input sensors at these loci, we next characterized NOT gates. For this, one of 9 different repressors was integrated at Int. 2 under control of P_tet_, together with its corresponding output promoter driving ymNeongreen, also at Int. 2. TetR itself remained integrated at Int. 1 as part of the sensor array. For every repressor, two or more output promoter variants were tested. One of the tested NOT gates did not show proper switching behavior (bm3R, data not shown).

**Figure 4:**
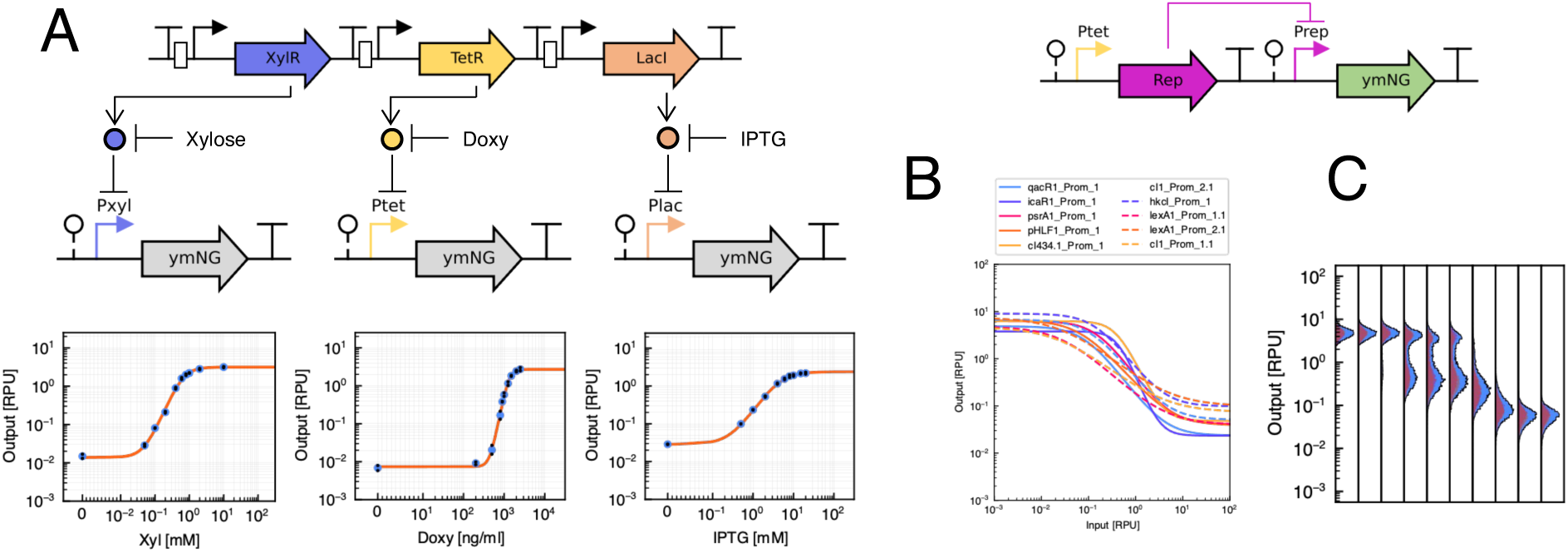
*A)* Measurement of the constitutive Ppfy1 promoter. The fluorescene values were used for the conversion of arbitrary fluorescence units into RPU. The measurement shows the mean of the median of a flow cytometry measurements performed twice on different days. *B)* The three sensors which are used as input gates. The sensors were inserted into the genome and measured with flow cytometry. The values show the mean of the distribution’s median performed on two different days. *C)* Not gate performance. The different repressors were under control of the Ptet promoter. The corresponding promotor of each repressor is controlling ymNeongreen. The response function are shown in RPU and represent the mean of the distribution’s median performed on two different days.

### 2.4 NOR gates realize two-input logic

Finally, we tested whether the NOR gates function as expected. NOR gates were built by adding a second copy of the repressor, under control of P_xyl_, at Int. 3, so that both repressor copies converge on the same output promoter driving ymNeongreen at Int. 2 (Figure 5 a). To integrate this second copy without triggering homologous recombination with the existing copy at Int. 2, its codon usage was altered while preserving the encoded protein sequence. ymNeongreen expression is expected only when both inputs are absent. Addition of either inducer alone, or both together, should repress output, realizing NOR logic behavior. Here, we present one representative NOR gate as proof of principle, confirming that the design functions as intended (Figure 5b) Characterization of the remaining NOR gate combinations can be found in the Supplementary Information (Supplementary Figure 3).

**Figure 5:**
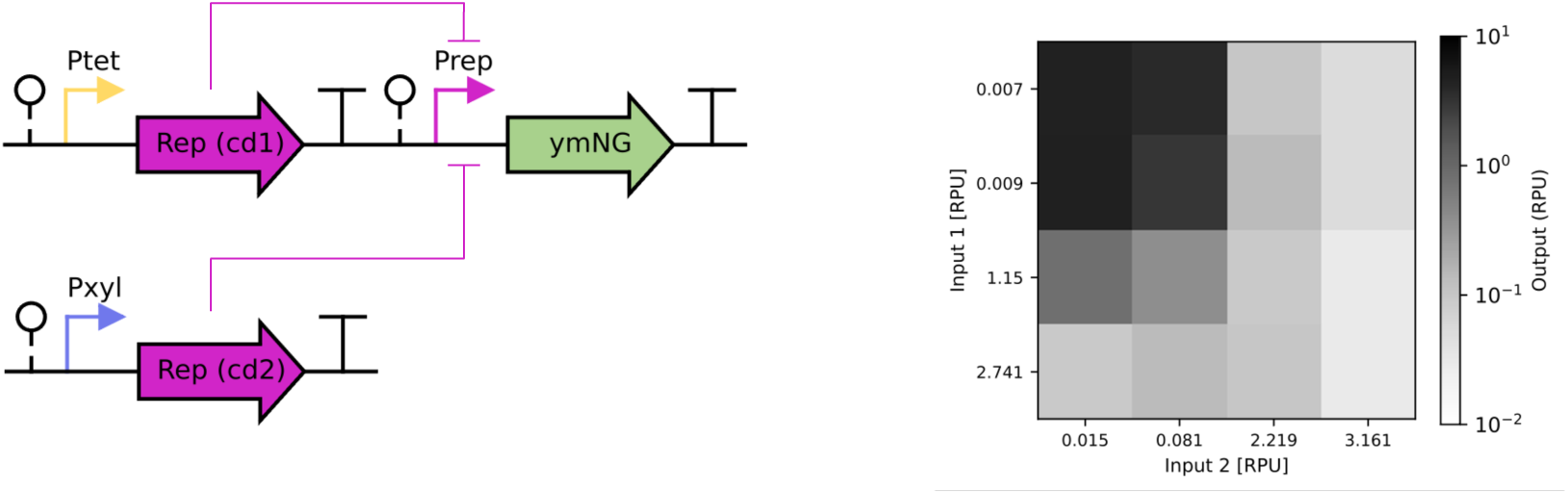
Response function of the NOR gates. NOR gates consist of the same repressor controlled by P_tet_ and P_xyl_ independently. Repressors have different codon usage to suppress homologous recombination. Response of qacR gate is shown. Input and output is shown as RPU.

## 3 Summary

We have developed a toolkit that bridges the Cello library for yeast with the widely used cloning scheme of the Yeast Toolkit (YTK). Everything presented in this work is therefore fully compatible with both YTK and its extension, the Multiplexed Yeast Toolkit (MYT). We provide the NOT and NOR gates from the Cello library fully characterized within the YTK cloning framework. This gives users the ability to build synthetic circuits using parts already established in their labs, simplifying exchange and comparability between groups. Of the nine repressors tested, eight showed the expected switching behavior. One gate (bm3R) did not display proper repression under our conditions, consistent with the AAGA overhang altering Kozak-driven expression levels, as observed in the Kozak variant screen (Figure 3). To further streamline assembly and make use of the insulators from the Cello library, we equipped 40 Level 1 backbones with defined orders of insulators. For the backbone in which we replaced the original RiboJ60 ribozyme with a VtmoJ ribozyme to avoid interference with the BsmBI/BsaI reactions, we demonstrated that the insulator blocks transcriptional read-through from an upstream transcriptional unit. These backbones were designed for three integration loci, but can be adapted to other loci if needed, and remain compatible with existing parts and backbones from previous toolkits. We also subdivided Type 2 parts into smaller subparts, giving users more freedom in 5’ UTR design: the T at position -1, previously fixed by the TATG overhang between Type 2 and Type 3 parts, is no longer constrained. As an example, we screened 9 different 10 bp Kozak sequences. In the same way, users can substitute different Kozak sequences or design longer 5’ UTRs depending on their needs. Altogether, YLT extends existing, widely used MoClo-based yeast toolkits with highly characterized parts and detailed usage guidelines, helping researchers speed up the assembly of complex genetic circuits. To support this, we also provide an automated data processing pipeline for gating and RPU conversion of flow cytometry data, adapted from the ColiToolkit (CTK) (*22*) framework and tailored with a new gating procedure for use with *S. cerevisiae*, reducing manual effort in future characterization efforts. Combined with automation platforms, this workflow could be scaled further to increase throughput for larger-scale circuit design efforts.

## 4 Materials and Methods

### 4.1 Strains and growth media

*S. cerevisiae* BY4741 (*MATa his3*Δ*1 leu2*Δ*0 met15*Δ*0 ura3*Δ*0)* were used in all experiments. For auxotrophic selection, synthetic defined agar (SDA) and the appropriate complete supplement mixture (CSM) lacking the auxotrophic markers was used (Sunrise Science, https://sunrisescience.com, CSM-U, CSM-UH, CSM-LH, CSM-UHL). For liquid cultures, Synthetic defined medium (SD) mixed with CSM lacking the appropriate auxotrophic markers were used (Sunrise science, https://sunrisescience.com, CSM, CSM-UH, CSM-LH, CSM-UHL).

For Doxycycline induction experiments, we used doxycycline (doxy) (Sigma Aldrich, https://www.sigmaaldrich.com/DE/de). Doxy was dissolved in water and stored at -20 degree in light-safe tubes for long term storage. Fresh master stocks were prepared prior to the induction experiment in water and stored at -20 degree for short term storage.

For xylose induction experimentsm, xylose (Carl Roth, https://www.carlroth.com/) was dissolved in water and stored at -20 degree for long term storage. Fresh master stocks were prepared prior to the induction experiment in water and stored at 4 degree for short term storage.

For IPTG induction experiments, IPTG (Thermofisher, https://www.thermofisher.com) was dissolved in water and stored at -20 degree for long term storage. Fresh master stocks were prepared prior to the experiments in water and stored at -20 degree.

Cloning of plasmids was performed using NEB 10-beta or NEB 5-alpha cells https://www.neb-online.de/shop_d. Transformed cells were selected on Lysogeny Broth (LB) plates with the appropriate antibiotic (chloramphenicol, ampicillin, or kanamycin)

### 4.2 Cloning of YLT plasmids

The cloning of the lvl 0 plasmids was done by ordering the different parts at Twist (https://www.twistbioscience.com) with the appropriate overhangs depending on the part type and insertion into pYTK001 (*4*). The sequences for the insulator, promoter and repressors were adapted from (*14*) and altered if it had an internal BsmBI, BsaI or NotI sites. Type 1 parts were cloned by gibson assembly (*23*) into BsmBI digested pYTK001, due to its internal BsmBI sites. Pre-assembled backbones were cloned by inserting the correct type 1 and 5 parts together with a spacer (pYTK047) into pYT095. The Spacer was exchanged by a GFP dropout from pYTK047 via Gibson cloning. Golden Gate cloning steps were performed following the instructions from (*4*).

### 4.3 Flow Cytometry Data Preprocessing and Cleaning

The gating and data analysis of the flow cytometry data was performed using a Python Notebook (see Data Availability Statement and GitHub repository). The Notebook is based on the gating and analysis pipeline of the ColiToolkit (CTK) by Mejlsted et al. (*22*) and adapted to the needs of the YLT. Major modifications include a new gating procedure that has been tailored to S. cerevisiae. Data cleaning (outlier detection) and RPU conversion follow the methods described in (*22*) and are restated here for methodological completeness.

The flow cytometry data is gated using the FlowCal library (*24*) and follows a two gate approach. The first gate performs density gating on the height of forward and side scatter and preservers 95% of the cell events. The second gate is defined on the forward scatter area and height and removes non single cell events. Details on the selection of the single cell gate can be found in the Supplementary Information. The gates are calibrated per replicate on the negative control. Per replicate gates are then applied to cytometry data processed.

Data cleaning and RPU calculation is as described by Mejlsted et al. (*22*) and repeated here for completeness. For data cleaning (outlier detection), we only consider experimental conditions with small deviations between replicates. In particular, we consider the pairwise ratios of the replicates’ median values and pool the data only in case all ratios are smaller than or equal to eight. Otherwise, we discard all three replicates for the respective experimental conditions (a particular inducer concentration). In addition, we consider only replicates with at least 1000 cell events. For later model calibration, we preferred to not introduce any bias and instead discard the experimental condition as a whole as the model curvature can be inferred from neighboring experimental conditions. The ratio of eight proved robust for discarding high deviation replicates and tolerant to natural deviations. Both procedures were chosen to ensure robust data cleaning.

Further preprocessing includes conversion to relative promoter units (RPUs) to quantify the relative promoter activity in comparison to the reference promoter Ppfy1 (*14*). The conversion factor *γ* to convert raw fluorescence intensity values into RPU was determined by using the following formula (*13*).

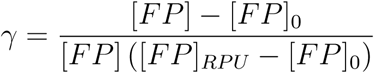

Here, [*FP*] is the median fluorescence of the sample, [*FP*]_0_ is the median fluorescence of the autofluorescence of the control (pooled negative controls), and [*FP*]*_RPU_* is the median fluorescence of the cells containing the reference plasmid with the reference promoter Ppfy1 (*14*). All raw fluorescence values were then rescaled by multiplication with *γ* to yield RPU values.

### 4.4 Model Calibration

The model calibration of the Yeast Logic Toolkit follows the model calibration by Mejlsted et al. (*22*). For reasons of completeness, we here restate the definition of the models, the priors, the likelihood as well as the description of the parallel tempering (*25*, *26*) algorithm and the hyperparameters used.

To represent the dose-response curves of input sensors and gates analytically, we calibrated models to the median RPU dose-response. The model calibration uses the data set *D* = {*x_i_, y_i_*}. Here, the *x_i_* ∈ ℝ*_≥_*_0_ either represent the inducer concentrations in case of the input sensors or the corresponding input sensor’s median output in RPU, which is the gate’s input. In both cases, the *y_i_* ∈ ℝ*_>_*_0_ are the corresponding median outputs in RPU. To model the dose-response curves, we use the activatory Hill equation defined as

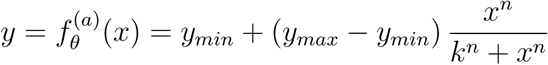

for the input sensors, and the inhibitory Hill equation

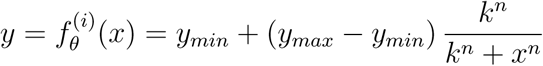

for the gates. Parameters *y_min_*, *y_max_*, and *k* are in RPU and define the dynamic range of the output (*y_min_* and *y_max_*) as well as the location of the transition region (*k*). *n* is the Hill coefficient and defines the steepness and, in turn, the length of the transition region. *θ* = (*y_max_, y_min_, n, k*) represents the model’s parametrization.

The algorithm employed for model calibration is parallel tempering (*25*, *26*), a Markov chain Monte Carlo algorithm. Parallel tempering uses Markov chains at different temperatures to draw samples from the posterior distribution. This allows one to explore multimodal distributions through sample exchange between chains, and was successfully applied to model calibration of chemical reaction networks previously (*27*). As the experimental values as well as the parameters *θ* span multiple orders of magnitude, we will consider in both cases the logarithmic domain for the calculation of differences. For the experimental values, this ensures that the model’s deviation to the data is treated in dependence to the order of magnitude, while in case of the parameters, the algorithmic behavior is improved.

The posterior distribution *p*(*θ* | *D*) over the parameters *θ* in dependence to the median dose-response data *D* is defined as

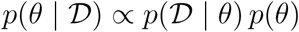

with prior *p*(*θ*) and likelihood *p*(*D* | *θ*). The prior encodes our initial assumptions on the parameters. As we assume the parameters to be independent, the prior factorizes into

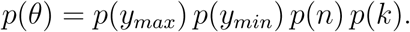

Further, we assume that the probability of *y_max_* and *y_min_* is highest in the range [*ŷ_max_,* 2 *ŷ_max_*] and [0.5 *ŷ_min_, ŷ_min_*] respectively, where *ŷ_max_* = max*_i_ y_i_* and *ŷ_min_* = min*_i_ y_i_*. We encode this as

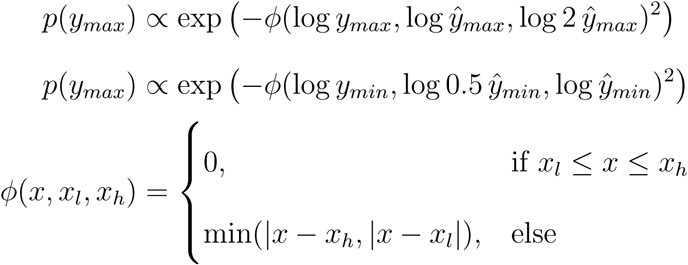

and we set *p*(*n*) = *p*(*k*) = 1, encoding no further assumptions. Please note that defining the distributions in terms of proportionalities is sufficient only for estimating the posterior distribution, respectively the maximum *a posteriori* (MAP) estimate, by parallel tempering or a Metropolis-Hastings algorithm.

The likelihood *p*(*D* | *θ*) characterizes how well the data matches the model with parameters *θ*. As both models (*f_θ_*^(*a*)^ and *f_θ_*^(*i*)^) have four parameters each, they can be treated identically wherefore we introduce *f_θ_* representing either of *f_θ_*^(*a*)^ and *f_θ_*^(*i*)^. We assume that the *y_i_* and *y_j_* are independent for *i* ≠ *j* and therefore define the factorized likelihood.

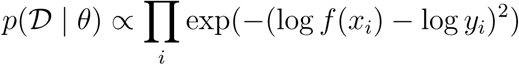

The calibrated parameter configuration 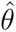 is then defined as the maximum *a posteriori* estimate.

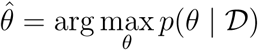

We identify 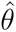 by sampling from the posterior distribution with parallel tempering. In particular, we define the initial parameter configuration to be

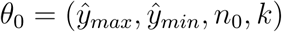

with *n*_0_ = 2 and *k*_0_ = 1 in the case of input sensor calibration or *k*_0_ = 0.01 in the case of gate calibration. Parallel tempering is executed for 10,000 steps with 10 walkers, each featuring 10 chains at different temperatures. From the 10^7^ posterior evaluations, we select 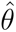 as the one maximizing the posterior. This process is performed independently for each input sensor and each gate.

The full model calibration pipeline is part of the Python Notebook for data processing (see Data Availability Statement).

## Supporting information

Plasmid Files

## 5 Data Availability Statement

All YLT plasmids (Supplementary Table 1) are available from the attached .zip file and will be made available on Addgene. Additional cloned plasmids are available upon request. The data and code is available at: https://github.com/Self-Organizing-Systems-TU-Darmstadt/YeastLogicToolkit.

## 6 Author Contributions

M.M. conceptualized the project, performed all experiments, and wrote the manuscript draft. E.K. provided the flow cytometry analysis, developed the model calibration pipeline and contributed to figure creation. H.K. conceptualized and supervised the project. All authors revised the manuscript.

## 7 Acknowledgements

The authors thank Jacob Mejlsted for fruitful discussions and Serin Joby Parekkadan for assistance with experimental preparation and execution. The work was supported by the Hessian Ministry of Science and Research, Arts and Culture as part of a LOEWE Top Professorship for H.K.

