## Supplemental Material for "A Yeast Logic Toolkit for Fast and Efficient Assembly of Genetic Circuits"

Table S1: List of toolkit plasmids

| Plasmid | Type | Description | *E. coli* marker |
|---|---|---|---|
| pYLTK001 | Type 1 | ConAS | CamR |
| pYLTK002 | Type 1 | ConA1 | CamR |
| pYLTK003 | Type 1 | ConA2 | CamR |
| pYLTK004 | Type 1 | ConA3 | CamR |
| pYLTK005 | Type 1 | ConA4 | CamR |
| pYLTK006 | Type 1 | ConA5 | CamR |
| pYLTK007 | Type 1 | ConA6 | CamR |
| pYLTK008 | Type 1 | ConA7 | CamR |
| pYLTK009 | Type 1 | ConBS | CamR |
| pYLTK010 | Type 1 | ConB1 | CamR |
| pYLTK011 | Type 1 | ConB2 | CamR |

Continued on next page

Table S1 – continued from previous page

| Plasmid | Type | Description | *E. coli* marker |
|---|---|---|---|
| pYLTK012 | Type 1 | ConB3 | CamR |
| pYLTK013 | Type 1 | ConB4 | CamR |
| pYLTK014 | Type 1 | ConB5 | CamR |
| pYLTK015 | Type 1 | ConB6 | CamR |
| pYLTK016 | Type 1 | ConB7 | CamR |
| pYLTK017 | Type 1 | ConCS | CamR |
| pYLTK018 | Type 1 | ConC1 | CamR |
| pYLTK019 | Type 1 | ConC2 | CamR |
| pYLTK020 | Type 1 | ConC3 | CamR |
| pYLTK021 | Type 2a | Ptet | CamR |
| pYLTK022 | Type 2a | Pxyl | CamR |
| pYLTK023 | Type 2a | Plac | CamR |
| pYLTK024 | Type 2a | PhkcI.1 | CamR |
| pYLTK025 | Type 2a | PhkcI.2 | CamR |
| pYLTK026 | Type 2a | PlexA1.1 | CamR |
| pYLTK027 | Type 2a | PlexA1.2 | CamR |
| pYLTK028 | Type 2a | PlexA2.1 | CamR |
| pYLTK029 | Type 2a | PlexA2.2 | CamR |
| pYLTK030 | Type 2a | PcI1.1 | CamR |
| pYLTK031 | Type 2a | PcI1.2 | CamR |
| pYLTK032 | Type 2a | PcI2.1 | CamR |
| pYLTK033 | Type 2a | PcI2.2 | CamR |
| pYLTK034 | Type 2a | Pbm3R1.1 | CamR |

Continued on next page

Table S1 – continued from previous page

| Plasmid | Type | Description | *E. coli* marker |
|---|---|---|---|
| pYLTK035 | Type 2a | Pbm3R1.2 | CamR |
| pYLTK036 | Type 2a | PcI434.1 | CamR |
| pYLTK037 | Type 2a | PcI434.2 | CamR |
| pYLTK038 | Type 2a | PicaR.1 | CamR |
| pYLTK039 | Type 2a | PicaR.2 | CamR |
| pYLTK040 | Type 2a | PphlF.1 | CamR |
| pYLTK041 | Type 2a | PphlF.2 | CamR |
| pYLTK042 | Type 2a | PqacR.1 | CamR |
| pYLTK043 | Type 2a | PqacR.2 | CamR |
| pYLTK044 | Type 2a | PpsrA.1 | CamR |
| pYLTK045 | Type 2a | PpsrA.2 | CamR |
| pYLTK046 | Type 2 | Ptet | CamR |
| pYLTK047 | Type 2 | Pxyl | CamR |
| pYLTK048 | Type 2 | Plac | CamR |
| pYLTK049 | Type 2 | PhkcI.1 | CamR |
| pYLTK050 | Type 2 | PhkcI.2 | CamR |
| pYLTK051 | Type 2 | PlexA1.1 | CamR |
| pYLTK052 | Type 2 | PlexA1.2 | CamR |
| pYLTK053 | Type 2 | PlexA2.1 | CamR |
| pYLTK054 | Type 2 | PlexA2.2 | CamR |
| pYLTK055 | Type 2 | PcI1.1 | CamR |
| pYLTK056 | Type 2 | PcI1.2 | CamR |
| pYLTK057 | Type 2 | PcI2.1 | CamR |

Continued on next page

Table S1 – continued from previous page

| Plasmid | Type | Description | *E. coli* marker |
|---|---|---|---|
| pYLTK058 | Type 2 | PcI2.2 | CamR |
| pYLTK059 | Type 2 | Pbm3R1.1 | CamR |
| pYLTK060 | Type 2 | Pbm3R1.2 | CamR |
| pYLTK061 | Type 2 | PcI434.1 | CamR |
| pYLTK062 | Type 2 | PcI434.2 | CamR |
| pYLTK063 | Type 2 | PicaR.1 | CamR |
| pYLTK064 | Type 2 | PicaR.2 | CamR |
| pYLTK065 | Type 2 | PphlF.1 | CamR |
| pYLTK066 | Type 2 | PphlF.2 | CamR |
| pYLTK067 | Type 2 | PqacR.1 | CamR |
| pYLTK068 | Type 2 | PqacR.2 | CamR |
| pYLTK069 | Type 2 | PpsrA.1 | CamR |
| pYLTK070 | Type 2 | PpsrA.2 | CamR |
| pYLTK071 | Type 2b3 | Kozak 4 - BM3R1.1 | CamR |
| pYLTK072 | Type 2b3 | Kozak 4 - BM3R1.2 | CamR |
| pYLTK073 | Type 2b3 | Kozak 12 - CI.1 | CamR |
| pYLTK074 | Type 2b3 | Kozak 12 - CI.2 | CamR |
| pYLTK075 | Type 2b3 | Kozak 15 - CI434.1 | CamR |
| pYLTK076 | Type 2b3 | Kozak 15 - CI434.2 | CamR |
| pYLTK077 | Type 2b3 | Kozak 34 - HKCI.1 | CamR |
| pYLTK078 | Type 2b3 | Kozak 34 - HKCI.2 | CamR |
| pYLTK079 | Type 2b3 | Kozak 8 - IcaR.1 | CamR |
| pYLTK080 | Type 2b3 | Kozak 8 - IcaR.2 | CamR |

Continued on next page

Table S1 – continued from previous page

| Plasmid | Type | Description | *E. coli* marker |
|---|---|---|---|
| pYLTK081 | Type 2b3 | Kozak 14 - LexA.1 | CamR |
| pYLTK082 | Type 2b3 | Kozak 14 - LexA.2 | CamR |
| pYLTK083 | Type 2b3 | Kozak 1 - PhlF.1 | CamR |
| pYLTK084 | Type 2b3 | Kozak 1 - PhlF.2 | CamR |
| pYLTK085 | Type 2b3 | Kozak 10 - PsrA.1 | CamR |
| pYLTK086 | Type 2b3 | Kozak 10 - PsrA.2 | CamR |
| pYLTK087 | Type 2b3 | Kozak 7 - QacR.1 | CamR |
| pYLTK088 | Type 2b3 | Kozak 7 - QacR.2 | CamR |
| pYLTK089 | Type 3 | BM3R1.1 | CamR |
| pYLTK090 | Type 3 | BM3R1.2 | CamR |
| pYLTK091 | Type 3 | CI.1 | CamR |
| pYLTK092 | Type 3 | CI.2 | CamR |
| pYLTK093 | Type 3 | CI434.1 | CamR |
| pYLTK094 | Type 3 | CI434.2 | CamR |
| pYLTK095 | Type 3 | HKCI.1 | CamR |
| pYLTK096 | Type 3 | HKCI.2 | CamR |
| pYLTK097 | Type 3 | IcaR.1 | CamR |
| pYLTK098 | Type 3 | IcaR.2 | CamR |
| pYLTK099 | Type 3 | LexA.1 | CamR |
| pYLTK100 | Type 3 | LexA.2 | CamR |
| pYLTK101 | Type 3 | PhlF.1 | CamR |
| pYLTK102 | Type 3 | PhlF.2 | CamR |
| pYLTK103 | Type 3 | PsrA.1 | CamR |

Continued on next page

Table S1 – continued from previous page

| Plasmid | Type | Description | *E. coli* marker |
|---|---|---|---|
| pYLTK104 | Type 3 | PsrA.2 | CamR |
| pYLTK105 | Type 3 | QacR.1 | CamR |
| pYLTK106 | Type 3 | QacR.2 | CamR |
| pYLTK107 | Type 3 | TetR | CamR |
| pYLTK108 | Type 3 | LacI | CamR |
| pYLTK109 | Type 3 | XylR | CamR |
| pYLTK110 | Type 56781 | Sensor 1 | AmpR |
| pYLTK111 | Type 56781 | Sensor 1e | AmpR |
| pYLTK112 | Type 56781 | Sensor 2 | AmpR |
| pYLTK113 | Type 56781 | Sensor 2e | AmpR |
| pYLTK114 | Type 56781 | Sensor 3 | AmpR |
| pYLTK115 | Type 56781 | Sensor 3e | AmpR |
| pYLTK116 | Type 56781 | Sensor 4 | AmpR |
| pYLTK117 | Type 56781 | Sensor 4e | AmpR |
| pYLTK118 | Type 56781 | LocusA 1 | AmpR |
| pYLTK119 | Type 56781 | LocusA 1e | AmpR |
| pYLTK120 | Type 56781 | LocusA 2 | AmpR |
| pYLTK121 | Type 56781 | LocusA 2e | AmpR |
| pYLTK122 | Type 56781 | LocusA 3 | AmpR |
| pYLTK123 | Type 56781 | LocusA 3e | AmpR |
| pYLTK124 | Type 56781 | LocusA 4 | AmpR |
| pYLTK125 | Type 56781 | LocusA 4e | AmpR |
| pYLTK126 | Type 56781 | LocusA 5 | AmpR |

Continued on next page

Table S1 – continued from previous page

| Plasmid | Type | Description | *E. coli* marker |
|---|---|---|---|
| pYLTK127 | Type 56781 | LocusA 5e | AmpR |
| pYLTK128 | Type 56781 | LocusA 6 | AmpR |
| pYLTK129 | Type 56781 | LocusA 6e | AmpR |
| pYLTK130 | Type 56781 | LocusA 7 | AmpR |
| pYLTK131 | Type 56781 | LocusA 7e | AmpR |
| pYLTK132 | Type 56781 | LocusA 8 | AmpR |
| pYLTK133 | Type 56781 | LocusA 8e | AmpR |
| pYLTK134 | Type 56781 | LocusB 1 | AmpR |
| pYLTK135 | Type 56781 | LocusB 1e | AmpR |
| pYLTK136 | Type 56781 | LocusB 2 | AmpR |
| pYLTK137 | Type 56781 | LocusB 2e | AmpR |
| pYLTK138 | Type 56781 | LocusB 3 | AmpR |
| pYLTK139 | Type 56781 | LocusB 3e | AmpR |
| pYLTK140 | Type 56781 | LocusB 4 | AmpR |
| pYLTK141 | Type 56781 | LocusB 4e | AmpR |
| pYLTK142 | Type 56781 | LocusB 5 | AmpR |
| pYLTK143 | Type 56781 | LocusB 5e | AmpR |
| pYLTK144 | Type 56781 | LocusB 6 | AmpR |
| pYLTK145 | Type 56781 | LocusB 6e | AmpR |
| pYLTK146 | Type 56781 | LocusB 7 | AmpR |
| pYLTK147 | Type 56781 | LocusB 7e | AmpR |
| pYLTK148 | Type 56781 | LocusB 8 | AmpR |
| pYLTK149 | Type 56781 | LocusB 8e | AmpR |

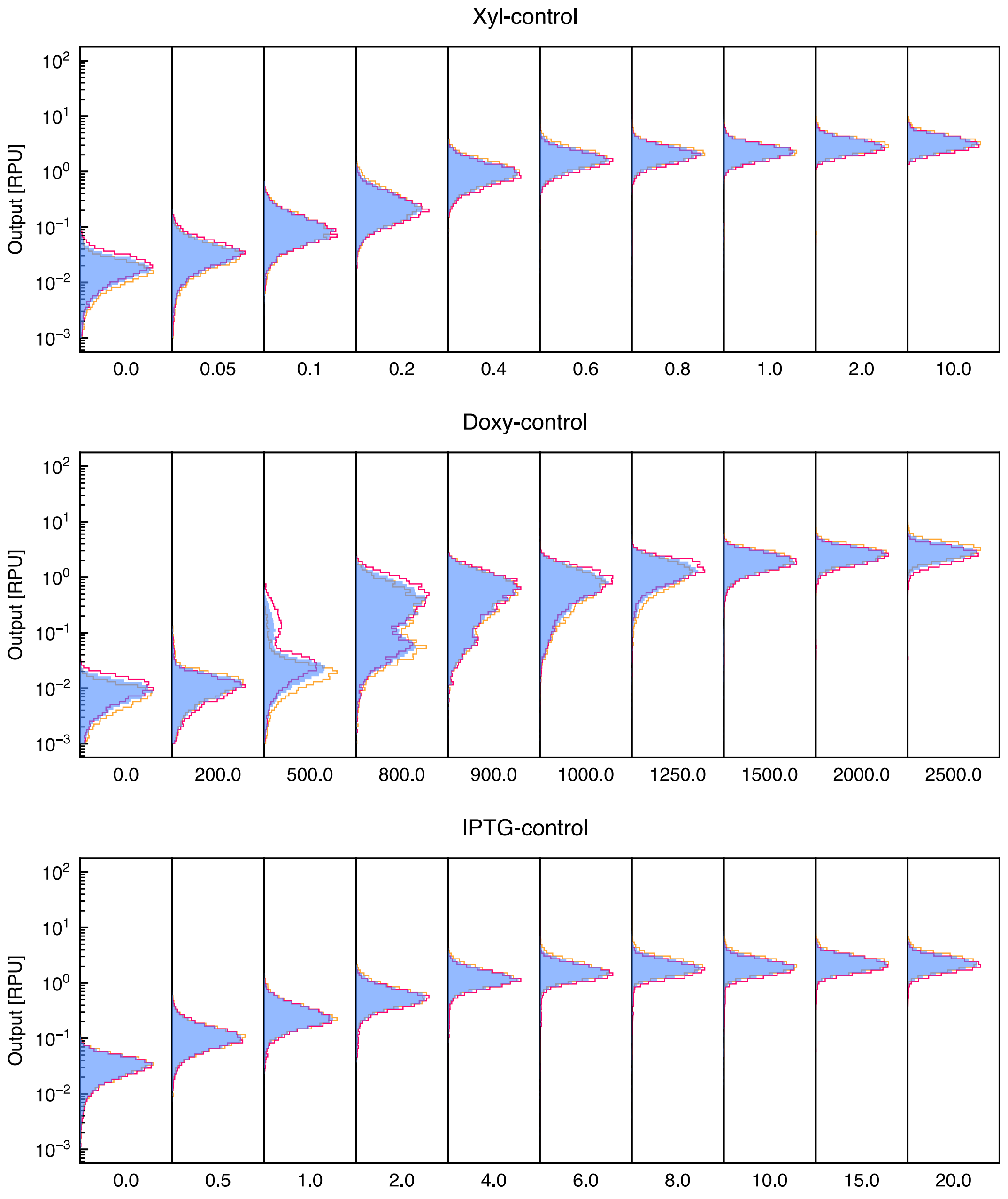


Figure S1: Individual populations of the three sensors across 10 different concentrations (mM for xylose, ng/ml for doxycycline, mM for IPTG). Individual populations can be seen in red and yellow, blue represents the merge of both.

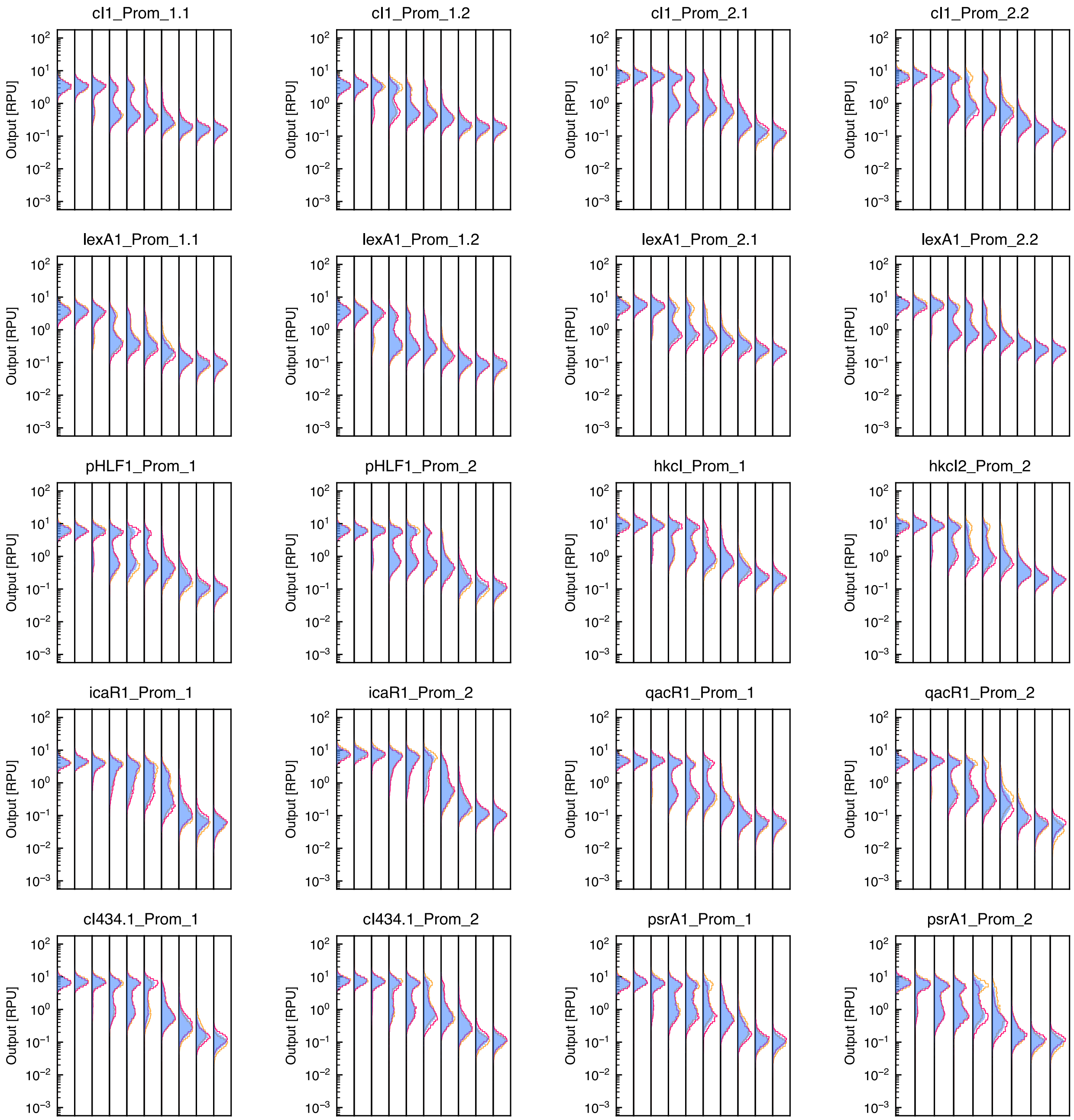


Figure S2: Individual populations of all NOT gates across 10 different RPU inputs. Individual populations can be seen in red and yellow, blue represents the merge of both.

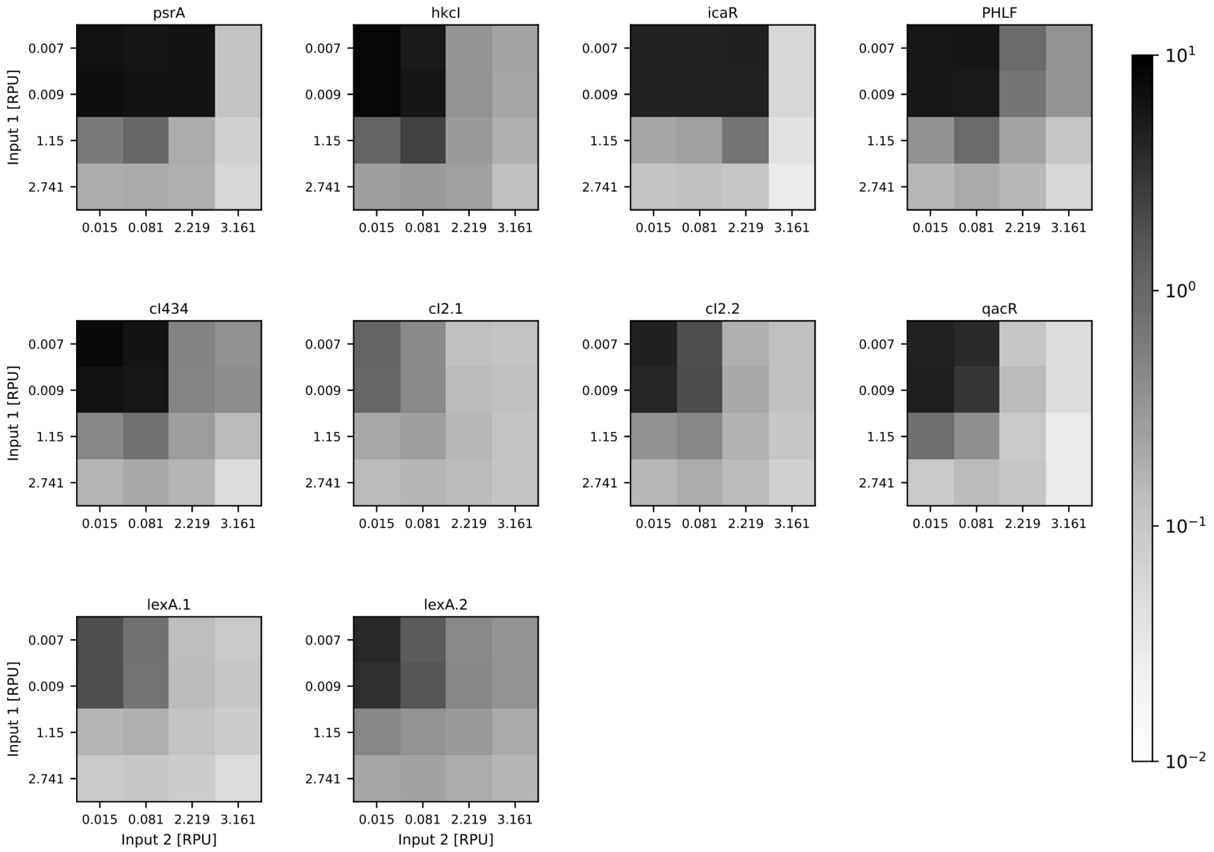


Figure S3: NOR gate RPU output across 16 different combinations of two inputs

### 1 Automated Analysis and Model Calibration Suite

Essential to the modular use of the parts in this toolkit is the analysis and model calibration, allowing for targeted part selection and *in silico* design. We provide our pipeline for analysis and model calibration, which we designed for both reproducibility but also reusability for new applications, as a Jupyter Notebook. The notebook is available at `https://github.com/Self-Organizing-Systems-TU-Darmstadt/YeastLogicToolkit`.

Here we focus on a tutorial to adapt the Notebook to your personal project and detail the gating procedure afterwards.

#### 1.1 Tutorial

To use the Notebook for your project, you can follow this section to adapt it to your needs. The Python Notebook can be separated into five parts: (I) setup, (II) data loading, (III) data conversion, (IV) model calibration, and (V) visualizations (see Figure S4).

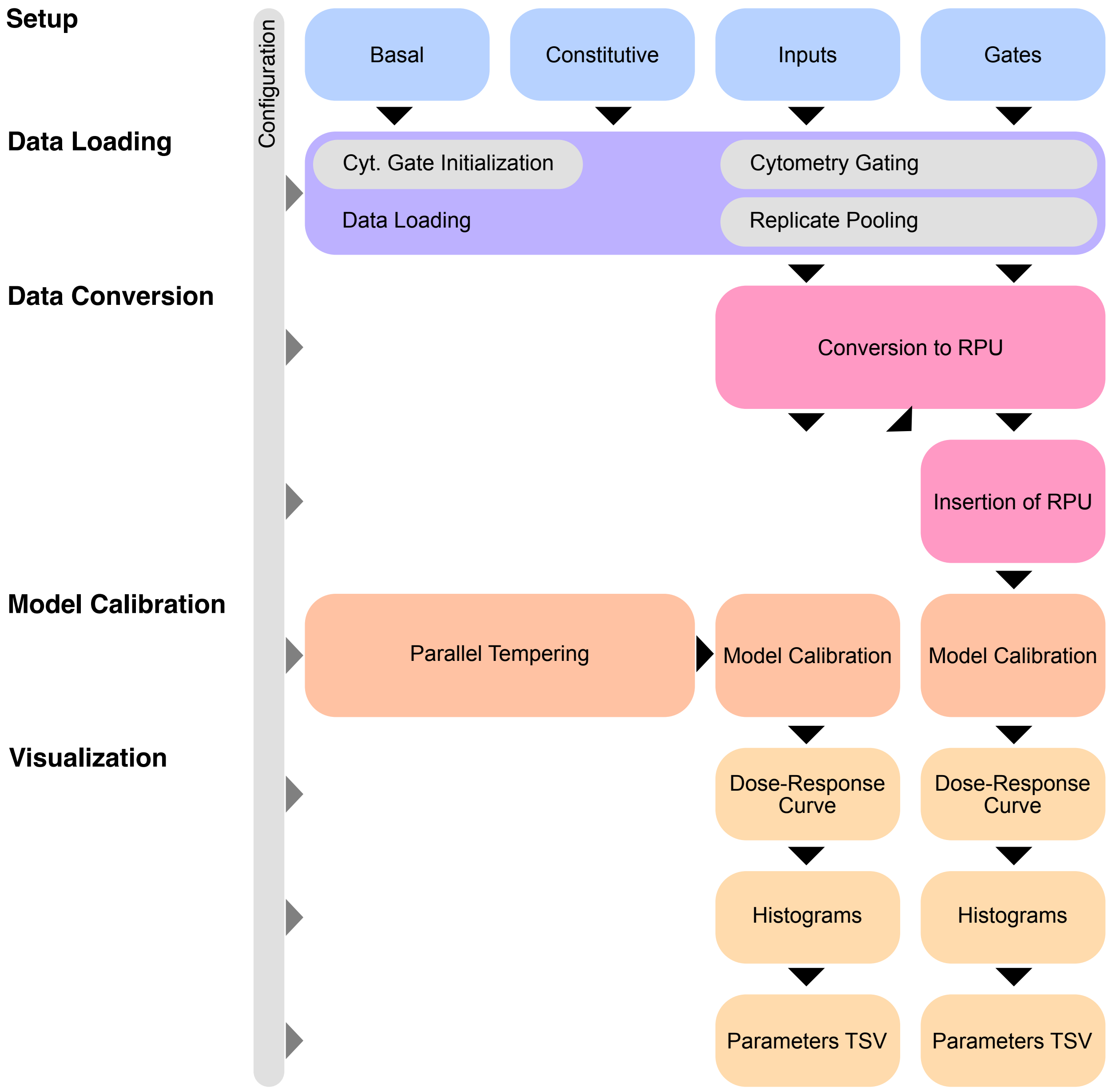


Figure S4: Overview on the cytometry data analysis and model calibration pipeline.

**(I) Setup** Here one defines the paths to the data, the units, the identifier of the reference promoter and the negative control, the input sensor used for gate characterization and the directory for storing the generated visualizations as well as some general setups for parallel tempering and visualizations.

```python
data_dir = pathlib.Path("data/")

data_dir_constitutive = data_dir / "constitutive/"
data_dir_inputs = data_dir / "inputs/"
data_dir_gates = data_dir / "gates/"
data_dir_basal = data_dir / "basal/"

✓ [2] < 10 ms
```

Figure S5: Code section where the input directories are specified. The directories have to adhere the structure as defined in (II) Data Loading.

```python
reference_promoter = "Positive"  # The constitutive reference promoter which serves as baseline for the RPU conversion
alternative_reference_promoter_name = {
    "Positive": "Positive"  # Currently, the reference promoter is part of the dataset in two different names
}
inducer_units = {
    "ind1": "ng-ml",
    "ind2": "mM",
    "ind3": "mM"
}
inducer_units_alias = {
    "ng-ml": "ng/ml",
    "mM": "mM",
}
inducer_molecule_type = {
    "Doxy-control": "Doxy",
    "Xyl-control": "Xyl",
    "IPTG-control": "IPTG"
}
controlling_input_sensor = "Doxy-control"  # The input sensor used for the characterization of the gate plasmids
facs_channel = "FL1-A"  # The fluorescence FACS channel we are interested in
✓ [4] < 10 ms
```

Figure S6: Code section where the inducer units, aliases, molecule types and input sensor for gate characterization as well as fluorescence channel is defined.

**(II) Data Loading** To use the script for ones own project, it is critical that the following directory structure is preserved.

```
data/
├── basal/
│   ├── replicate 1/
│   ├── replicate 2/
│   └── ...
└── constitutive/
    └── ...
```

```
│
├── gates/
│   └── ...
└── inputs/
    └── ...
```

In the Notebook, the path to the directories `basal`, `constitutive`, `gates`, and `inputs` is defined. Each of these directories includes one or more directories representing the individual replicates, where the corresponding `.fcs` files of all characterized parts are provided. To be compliant to the Notebook, each part needs to follow the naming convention `[NAME] [INDUCER LEVEL][INDUCER UNIT].fcs` (i.e. `cI1_Prom_1.1 200ng-ml.fcs` with `[NAME]=cI1_Prom_1.1`, `[INDUCER LEVEL]=200`, and `[INDUCER UNIT]=ng-ml`). From the directory structure and the part names, the Python Notebook will then automatically infer the replicate, the part name, the inducer level and the corresponding unit and cluster together data corresponding to the same part. You next need to add the proper part name for the reference promoter and choose the basal expression for the successful conversion to RPU. In case you like to add a new category, please make sure that you select the appropriate method to load your data. If the same part is present for different inducer levels, you should use `load_inducible_data( )`. Otherwise please use `load_constitutive_data( )`.

**(III) Data Conversion** During loading of the files, the Notebook will already gate the data based on the cytometry gates derived from the negative controls and merge replicates where appropriate. Afterwards follows the conversion to RPU as described in the Methods section of this work. After obtaining the output RPU levels for the input devices and the gates, we continue with inserting the output RPU levels of the inputs as input RPU levels to the gates. By that we obtain the RPU to RPU mapping for the gates, which we also will use for the model calibration.

**(IV) Model Calibration** During model calibration, we use parallel tempering (*1*, *2*) with priors and likelihoods as defined in the Methods section. By that, we sample from the posterior $p(\theta|\mathcal{D})$ and can obtain the parameter fit as the maximum *a posteriori* (MAP) estimate. The algorithm also reports the best parameter configuration obtained and the corresponding loss (sum over squared differences of log model values and log median values). We first do so for each of the input devices and afterwards for each gate device. The inputs are represented by an activatory Hill equation to represent the activating behavior of inducer chemicals while the gates are fit to an inhibitory Hill equation, representing the repressor behavior of the transcription factors. In case you have inputs that don't behave activating or want to change the models, you can do so in the sections `Fit Inputs` and `Fit Outputs` while we define the models in the section `Define Models`.

**(V) Visualizations** Following the preprocessing of the data in (II) Data Loading and (III) Data Conversion as well as obtaining the calibrated models in (IV) Model Calibration, the gathered data is visualized. For the inputs as well as the gates, two types of diagrams are created. The first visualizes the median expression in RPU in a scatterplot. Thereby, the individual replicates are highlighted as small black dots while the pooled data is represented by the blue dot. If data for an inducer concentration is discarded (replicate outliers detected), the coresponding data is shown grayed out. If a calibrated model is present, the corresponding dose-response curve is added to the plot. This plot allows to evaluate the quality of the model calibration and suitability for *in silico* design. The second plot created includes histograms for the pooled data as well as the individual replicates per inducer concentration. This plot allows the qualitative assessment of how well the replicates agree and how they contribute to the pooled data. Again, discarded data is shown grayed out.

The final plot visualizes the dose-response curves of the main promoters by their calibrated models in a single plot. This allows comparison with respect to dynamic range and switching characteristics. In addition, we output `.tsv` files for the gates and inputs including the parameters identified by model calibration.

### References


1. Swendsen, R. H., and Wang, J.-S. Replica Monte Carlo Simulation of Spin-Glasses. *57*, 2607–2609.

2. Earl, D. J., and Deem, M. W. Parallel Tempering: Theory, Applications, and New Perspectives. *7*, 3910–3916.
